# HigH-ratiO partiaL proteolysiS with carriER proteome (HOLSER) Enables Global Structure Profiling and Site-resolved Elucidation of Ligand-Protein Interactions

**DOI:** 10.1101/2025.07.11.664381

**Authors:** Xuepei Zhang, Bohdana Sokolova, Zhaowei Meng, Hassan Gharibi, Hezheng Lyu, Amir Ata Saei, Massimiliano Gaetani, Roman A. Zubarev

**Author notes:** Correspondence and requests for materials should be addressed to M.G. and R.A.Z.;).

## Abstract

Understanding how cellular proteins interact with their environment, including endogenous and exogeneous molecules, is critical for elucidating mechanisms of cellular regulation and drug action. Partial proteolysis-based techniques offer peptide-level resolution of ligand-induced conformational changes but are limited by modest proteome coverage and depth, as well as sensitivity to the experimental conditions. To overcome these limitations, we developed higH ratiO partiaL proteolysiS with carriER proteome (HOLSER), an efficient workflow that features extended digestion time for reduced peptide yield variability as well as tandem mass tag multiplexing that includes full digests for enhanced proteome depth and sequence coverage as well as higher precision of peptide abundance measurements. We demonstrate HOLSER capabilities of probing structural changes on the scale of specific binding sites for kinase target mapping, individual protein domains for structural mapping of the FKBP-mTOR complex in response to rapamycin as well as global proteome structure profiling.

## INTRODUCTION

Characterizing protein native structures and their changes induced by protein-ligand interactions is essential for understanding cellular signaling, the molecular basis of disease and drug mechanisms. Traditional approaches such as thermal shift assays^1, 2^ and affinity-based pull-downs^3-5^ often lack the resolution or throughput required to profile complex proteomes under physiologically relevant conditions. In recent years, limited proteolysis-mass spectrometry (LiP-MS)^6-8^ and peptide-centric local stability assay (PELSA)^9^ have emerged as powerful tools to detect ligand-induced conformational changes at the peptide level. These profiling techniques rely on limited proteolysis of native proteins, where ligand-stabilized regions resist enzymatic cleavage, with detection of the ligand-induced changes via quantitative mass spectrometry.

The most recent such approach, PELSA^9^, uses lower-grade trypsin at a high enzyme-substrate ratio (1:2) in combination with a very short digestion time. PELSA opened way for scalable structural analysis at the proteome level and demonstrated spectacular results in selected cases. But despite these significant promises, the technique possesses some critical limitations. Due to the shallow digestion, PELSA exhibits modest proteome depth and limited sequence coverage, which significantly hampers the statistical power and generality of the analysis. Due to the short, one-minute digestion time, the approach may suffer from stochastic quantification variability, particularly for low-abundance proteins and transient interactions. This is exacerbated by the critical for drug target elucidation transition from the peptides to the protein level hinging in PELSA on the detection and correct quantification of a single “best” peptide. These shortcomings limit the applicability of this otherwise highly promising technique in high-throughput ligand screening and domain-level interaction mapping. The recently developed high-throughput PELSA (HT-PELSA), a scaled-up adaptation of the original PELSA approach, extends its analytical capabilities. However, the inherent limitation of low sequence coverage due to restricted proteolysis remains unaddressed^10^.

To address these challenges, we developed HOLSER – an optimized workflow for ligand-induced stability profiling in cell lysate under native conditions (Fig. 1A). Inspired by PELSA to use high enzyme-to-substrate (E/S) ratio of 1:1 (wt/wt) and lower-grade trypsin, HOLSER features several critical differences. First, the digestion time is extended from one to four minutes to improve cleavage uniformity and reduce stochastic variability of the peptide yield. Then, instead of the label-free analysis that suffers significantly from the random fluctuations of electrospray current^11^, HOLSER employs the more precise tandem mass tag (TMT)-based quantification. Also, data-dependent acquisition is used that is less prone to false positive results^12, 13^ than the data-independent acquisition (DIA) employed in PELSA. Importantly, in the same TMT set partial digests in HOLSER are multiplexed with “carrier” channels containing a digest obtained in full (8 h long) proteolysis. As in single-cell proteomics^14^, the main purpose of the carrier channel is to boost sensitivity and detection probability, especially for low-abundant peptides from protected regions, and thus increase the analysis depth (number of quantified peptides and proteins) and average protein sequence coverage. Also, comparison of the peptide abundances in the partial digest versus full digest provides the “accessibility factor” that reflects protein tertiary structure even in the absence of a ligand. With TMT, better precision of abundance measurements allowed us to reduce the number of biological replicates from 4 to 3 without statistical power loss, with an overall win in the instrumental time and a potential to add to the standard TMT-18 set three more samples. Finally, instead of a single “best” peptide, HOLSER use several peptides for determining the target protein, with their p-values combined using Fisher’s formula. This provides robust and statistically viable target identification.

**Figure 1.**
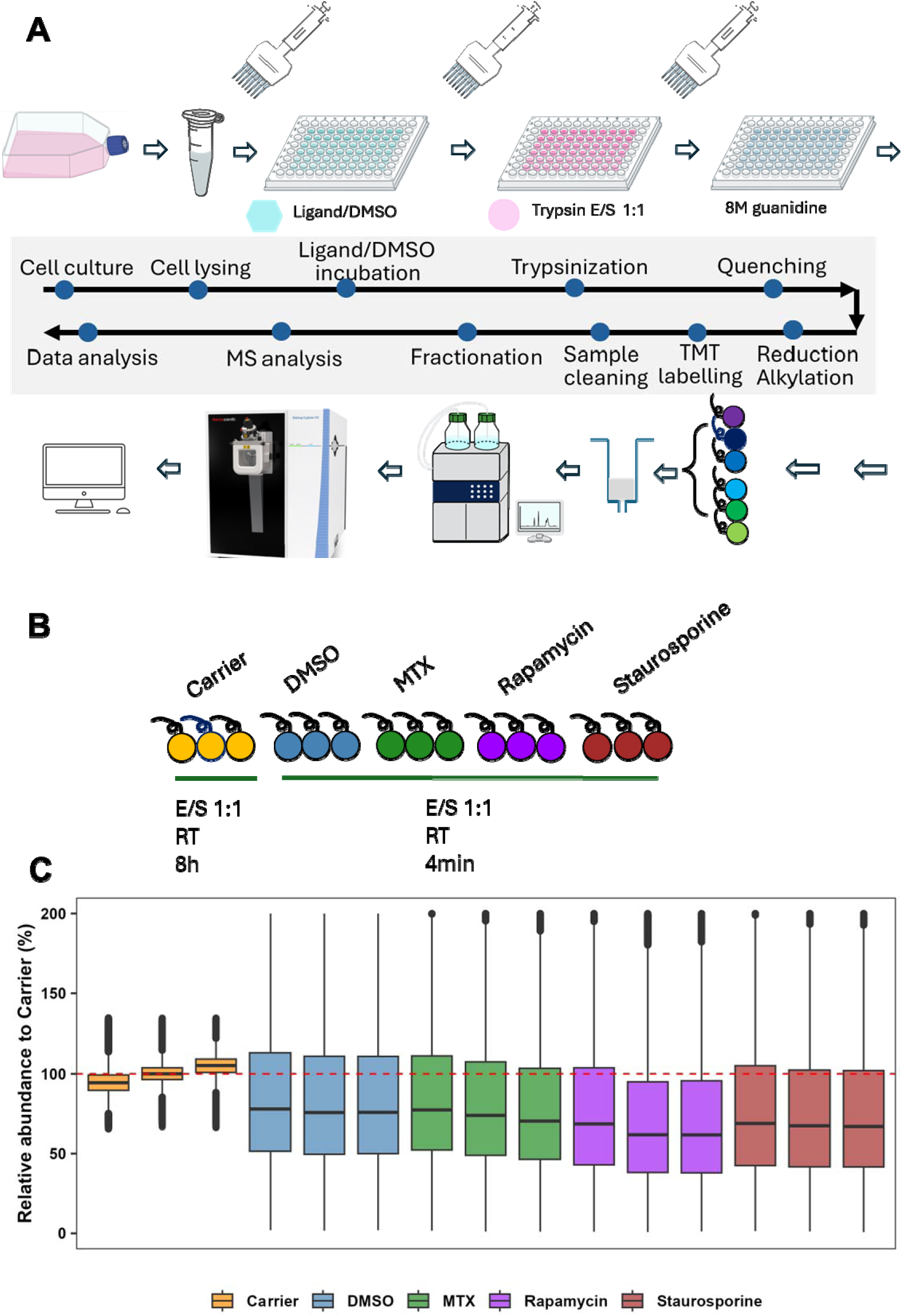
Overview of Experimental Workflow and TMT Labeling. **A,** Workflow of HOLSER. E/S ratio (wt/wt). **B,** TMT labeling scheme showing the assignment of unique tags to samples from different experimental groups and the inclusion of Carrier channels. **C,** Box plot showing relative peptide abundance for each TMT channel normalized to the carrier (dashed line at 100%)

In this proof of principle study, we apply HOLSER to verifying known ligand–protein interactions with high spatial resolution. We benchmark HOLSER performance against PELSA in global proteome profiling, staurosporine kinase target deconvolution, and targeted analysis of the FKBP–mTOR complex under rapamycin treatment. The results confirm that, as expected, HOLSER significantly improves proteome depth, protein sequence coverage, and the robustness of ligand-induced stabilization, additionally allowing for domain-level global structural mapping.

## RESULTS

### HOLSER enhances proteome analysis depth and sequence coverage

The 15 HeLa cell samples combined in a single TMT set included partial digests treated with vehicle DMSO (control), methotrexate (MTX), staurosporine and rapamycin, as well as full cell digest (Fig. 1B). The multiplexed TMT set was separated into 48 fractions by reversed-phase chromatography and analyzed by LC-MS/MS using an Orbitrap mass spectrometer (Thermo).

The obtained results were compared with data obtained in PELSA using HeLa cells, extracted in the form of final lists of proteins and peptides with relative quantification^9^. Protein sequence coverage of the proteins reported in HT-PELSA was calculated using the peptide-level results for identifying dasatinib targets in crude K562 cell lysates in that study^10^. For each protein, the start and end positions of all identified peptides were mapped onto the corresponding full-length protein sequence (retrieved from UniProt). Sequence coverage was defined as the percentage of amino acid residues of the protein that were covered by at least one peptide. Overlapping peptides were merged, and each protein is reported once. In HOLSER, the abundances in the TMT channels corresponding to partially digested samples were on average half of these in the fully digested “carrier” (Fig. 1C, Supplementary Table 1), reflecting the chosen balance between the depth of analysis and its specificity. At the peptide level, HOLSER quantified a total of 139,849 (Supplementary Table 2) peptides, compared to 71,708 peptides in PELSA – an increase by almost a factor of two. Surprisingly, only 60% of the PELSA peptides were found in our much deeper analysis. This startling result is in fact a frequent occurrence in proteomics^15^ and can be explained by both the sporadic nature of DDA as well as the enhanced false positive rate in DIA at low sequence coverage^16^. This result also warns against the reliance on a single peptide for target identification.

The protein-level comparison showed a larger (79%) overlap between the two techniques, but HOLSER could still not confirm the presence of 1465 PELSA proteins, while finding additional 3430 proteins. The average protein sequence coverage, 35% in HOLSER (Supplementary Table 1), was twice that of PELSA (17%) and HT-PELSA (17%), as would be expected for the inclusion in the TMT set of the full carrier digests. The *a priori* probability of finding in the dataset the correct target protein scales up respectively.

While shotgun proteomics can very rarely achieve 100% sequence coverage even for individual proteins, such as bovine serum albumin (BSA) frequently used for calibration and quality control^17^, obtaining ≥50% sequence coverage may be considered a desirable target. In that respect, PELSA – yielded 619 such proteins, while HOLSER - 2708 proteins (Supplementary Table 1), an increase by over 300%.

### HOLSER Reveals Peptide-Resolved Stabilization of DHFR Upon MTX Treatment

MTX is a favorite test molecule for novel drug target identification techniques, as its treatment significantly changes (increases) both abundance^18^ and solubility^19^ its cognate target, dihydrofolate reductase (DHFR). We have also successfully used MTX as a test molecule in our above-filter partial digestion technique AFDIP^20^, and it has also shown the expected behavior in PELSA. In HOLSER, multiple DHFR peptides exhibited strong ligand-induced protection (Fig. 2A). When peptides were ranked by a score combining the absolute fold change with statistical significance, two top candidates were DHFR-derived peptides (Fig. 2B, Supplementary Table 2).

**Figure 2.**
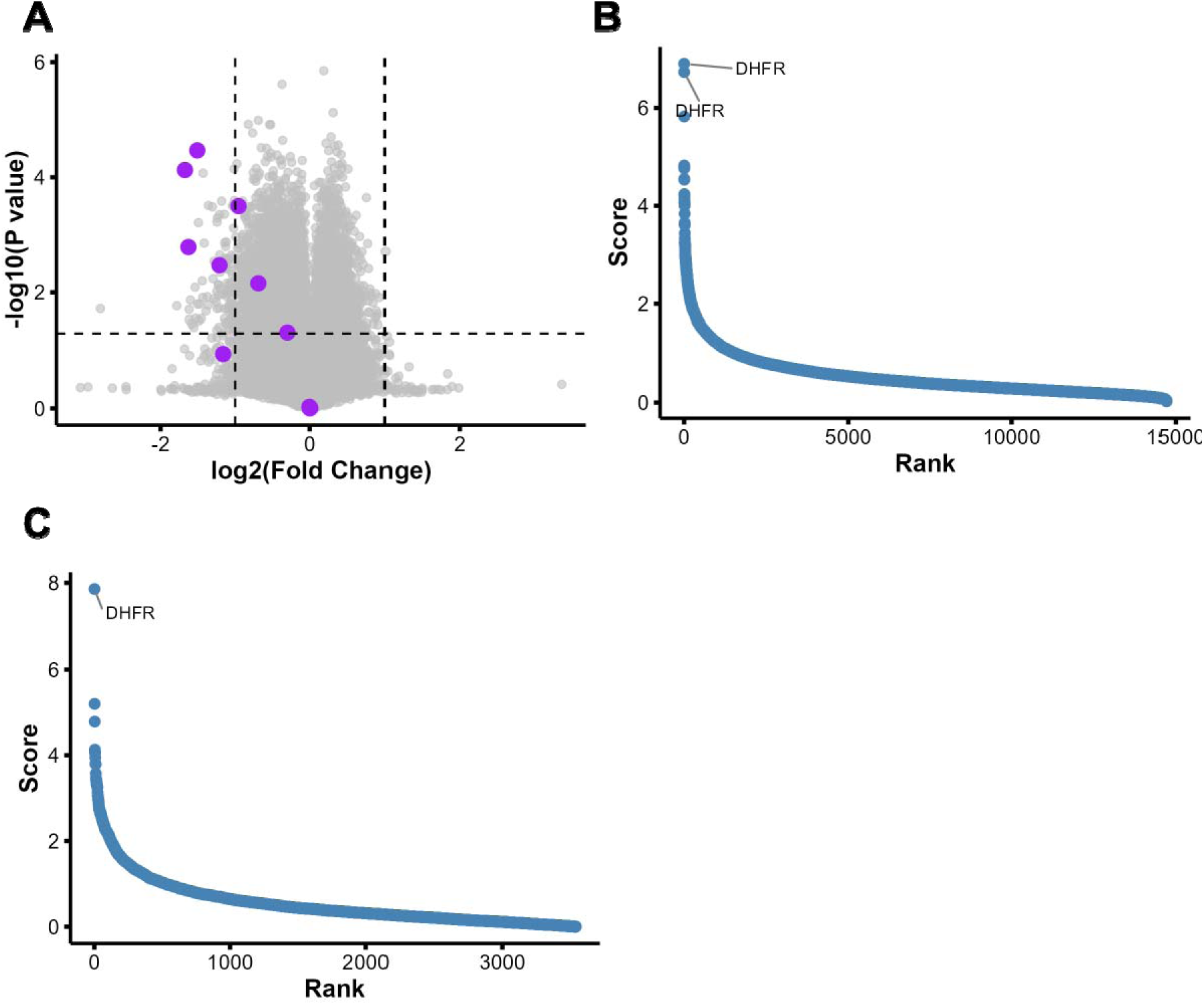
Peptide and protein abundance changes upon MTX treatment. **A,** Volcano plot displaying peptide abundance changes in MTX-treated samples compared with DMSO controls (n=3 biological replicates). Peptides derived from DHFR are highlighted in purple. Statistical significance was assessed by t-test. **B,** Ranked list of peptides according to their combined score (absolute log2 fold change × – log10 p_p_) calculated from replicate data. **C,** Ranked list of proteins according to their combined score (absolute log2 fold change × – log10 p value) calculated from replicate data.

Note that not all DHFR peptides have shifted strongly, and in AFDIP some peptides of the target protein shifted in the opposite direction. This is a distinct feature of the limited proteolysis techniques that provides structural resolution. Therefore, traditional for shotgun proteomics aggregation of the peptide data into protein results poorly suits these techniques. Here we combined the ranks of top three peptides for each protein to derive the protein rank (see Methods section), with DHFR expectedly appearing on the first position (Fig. 2C, Supplementary Table 3).

### HOLSER reveals targets of kinase-targeting staurosporine

Staurosporine is the broad-spectrum kinase inhibitor^21, 22^. Based on UniProt annotation, a total of 7932 peptides in the HOLSER dataset were confidently assigned to kinase proteins (Supplementary Table 2). Of these, 40.7% exhibited significant abundance changes, while in PELSA, only 22.0% of the 3643 kinase-derived peptides exhibited such changes. As a result, HOLSER identified 329 kinases as potential staurosporine targets, while PELSA - 157 kinases. The percentage of significant peptides per identified kinase was also higher, with median values of 37.5% in HOLSER compared to PELSA’s 18.2% (Fig. 3A). All 50 top-ranked peptides in HOLSER were derived from kinases. These results reveal better ability of HOLSER to identify target proteins. This is achieved at the expense of arguably lower than in PELSA specificity in determining the most likely site of ligand binding, provided that the low PELSA sequence coverage allows one to detect this peptide.

**Figure 3.**
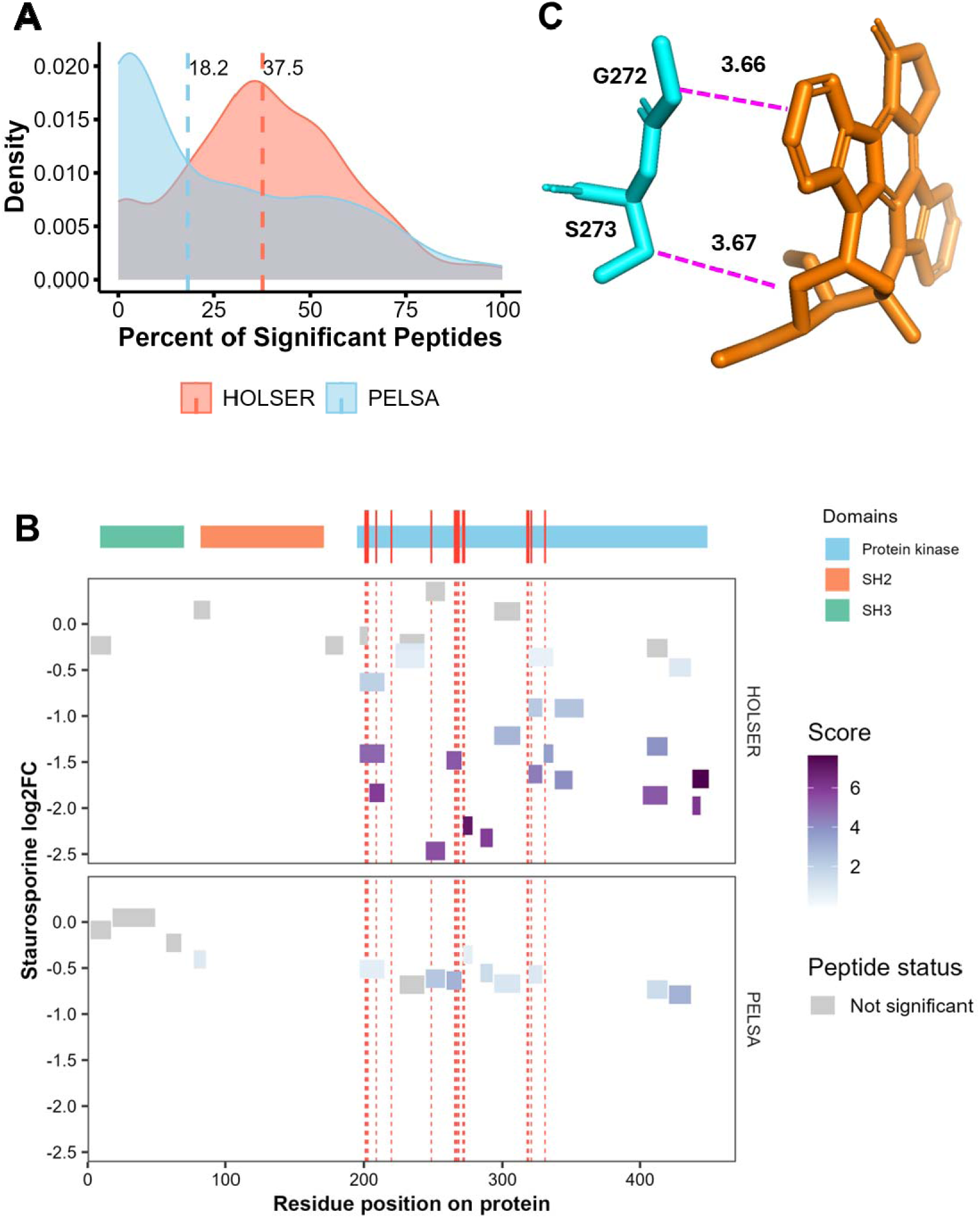
Kinase-targeting analysis using HOLSER and PELSA. **A,** Density plot showing the distribution of significant peptide percentages per kinase in HOLSER and PELSA. Dashed lines indicate the median values for each method. **B,** Mapping of peptides identified in HOLSER and PELSA onto the CSK sequence. **C,** Structural visualization of the CSK–staurosporine complex based on the crystal structure (PDB ID: 1BYG), rendered in PyMOL.

In CSK^23^, a well-characterized kinase target of staurosporine, HOLSER identified 29 peptides, while PELSA – 15 peptides. Of the HOLSER peptides, 25 (79%) showed significant changes, with 23 peptides mapping to the kinase domain, compared to 9 (60%) in PELSA. We used the known X-ray structure of the CSK-staurosporine complex^24^ (PDB ID: 1BYG) to correlate the peptide scores with the proximity to the binding site. Both in PELSA and in HOLSER, all higher-scoring peptides come from the kinase domain (Fig. 3B), but only in HOLSER the most-scoring peptides were close to the known staurosporine binding sites (Fig. 3B). The top-most peptide, GSLVDYLR, includes the G272 and S273 residues that both are just 3.7 Å away from the ligand (Fig. 3C, Supplementary Table 4). The two next-best peptides are adjacent to that peptide on its both sides (Fig. 3B).

One example where HOLSER with its deeper analysis uniquely detected a staurosporine target is DAKP1 with four peptides identified, of which the only significantly stabilized peptide covered kinase binding sites I160 and D161, while PELSA failed to detect any DAKP1 peptides (Supplementary Fig. 1A-B, Supplementary Table 5). In RPS6KA1, HOLSER identified 51 peptides (PELSA – 27). Among the HOLSER peptides, 32 (79%) showed significant changes, with 16 (50%) mapping to the kinase domain, compared to 6 (35%) peptides in PELSA. Also, in RPS6KA1 most significantly protected HOLSER peptides correctly identified the N-domain as the preferred staurosporine binding site^25^, while from PELSA data one could not make such conclusion with statistical certainty (Supplementary Fig. 1C).

We used the known X-ray structure of the RPS6KA1–staurosporine complex^24^ (PDB ID: 2Z7R) to correlate peptide scores with proximity to the binding site. One of the top-scoring peptides, LYLILDFLR, contains four residues (L141, D142, F143, L144) located 3.86 Å, 2.67 Å, 2.52 Å, and 3.36 Å from the ligand, respectively (Supplementary Fig. 1D, Supplementary Table 6). In PELSA, this peptide showed no significant difference (p-value > 0.05).

### HOLSER Profiling of FKBP-mTOR Complex Response to Rapamycin

Rapamycin targets FKBP proteins and mTOR, forming a ternary complex critical for mTOR pathway regulation^26, 27^. At the peptide level, 133 out of 439 detected peptides with belonging to FKBP and mTOR family (30%) showed in HOLSER statistically significant shifts, compared to 52 peptides out of 254 peptides (21%) in PELSA. Among the proteins, FKBP family members were clearly identified as the most likely target candidates (Fig. 4A, Supplementary Table 7), with FKBP3^28^ being the top candidate.

**Figure 4.**
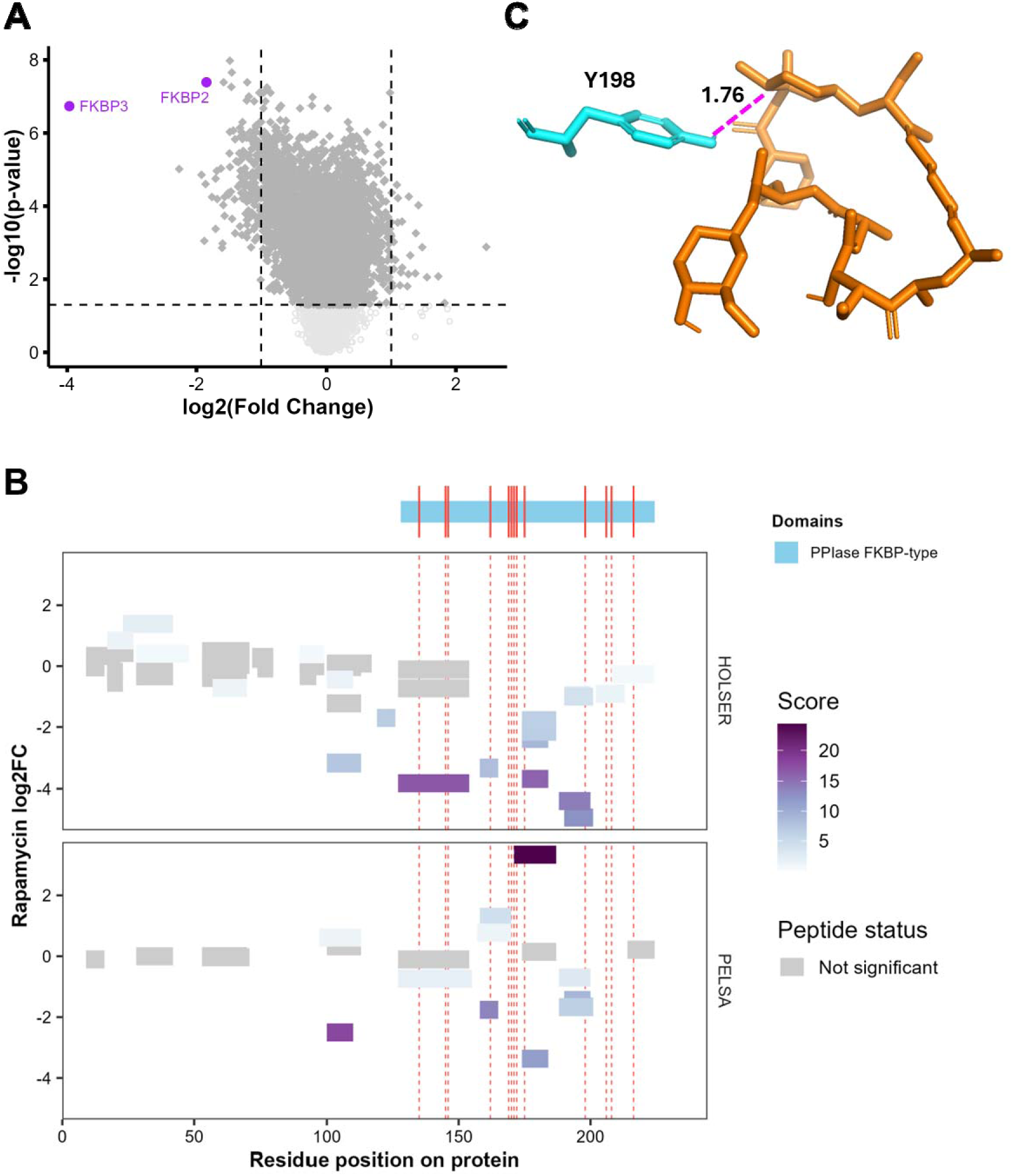
FKBP response to rapamycin treatment. **A**, Volcano plot displaying protein-level abundance changes in rapamycin-treated samples compared with DMSO controls (n = 3 biological replicates). **B,** Mapping of FKBP3-derived peptides identified in HOLSER and PELSA onto the protein sequence. **C,** Structural visualization of the FKBP3–substrate complex based on the crystal structure (PDB ID: 1PBK), rendered in PyMOL.

Sequence mapping of FKBP3-derived HOLSER peptides provided broader coverage across the protein sequence (46 peptides and 85.3 % sequence overage versus 20 and 64.3 %, respectively, in PELSA) and identified multiple additional peptides from the binding region (Fig. 4B). Notably, the most-protected in HOLSER peptide LEIEPEWAYGKK includes the residue Y198 located just 1.76 Å away from rapamycin in the FKBP3–rapamycin complex structure^29^ (PDB ID: 1PBK) (Fig. 4C, Supplementary Table 8). In PELSA, the peptide containing this residue has only a modest significance, with the most significant peptide from the binding region surprisingly showing reversed protection sign.

### HOLSER global proteome profiling

As already mentioned, the ratio between the peptide abundance in the partial digest of DMSO-treated channel and full digest provides the accessibility factor related to protein tertiary structure. As an example, Fig. 5A shows the HOLSER accessibility factors of KPNA2 protein and compares them the protection factors from HDX MS (Supplementary Table 9). A significant positive correlation (R = 0.42, p = 0.003) was observed between these two parameters. The FKBP3 protein exhibited an even better agreement (Fig. 5B, Supplementary Table 10), with a comparably high positive correlation (R = 0.46, p = 0.012). Comparisons of the protein structures with HDX MS protection factors against the HOLSER accessibility values (Fig. 5C and 5D) confirm the similarities, while the differences can at least partially be explained by the absence of heterogeneous protein-protein interactions and post-translational interactions in HDX MS that is performed on isolated, recombinant proteins as opposed to native lysate in HOLSER. Note that a single HOLSER analysis provides 10^5^ independent pieces of structural information on several thousands of proteins, making it one of the most informative structural approaches.

**Figure 5.**
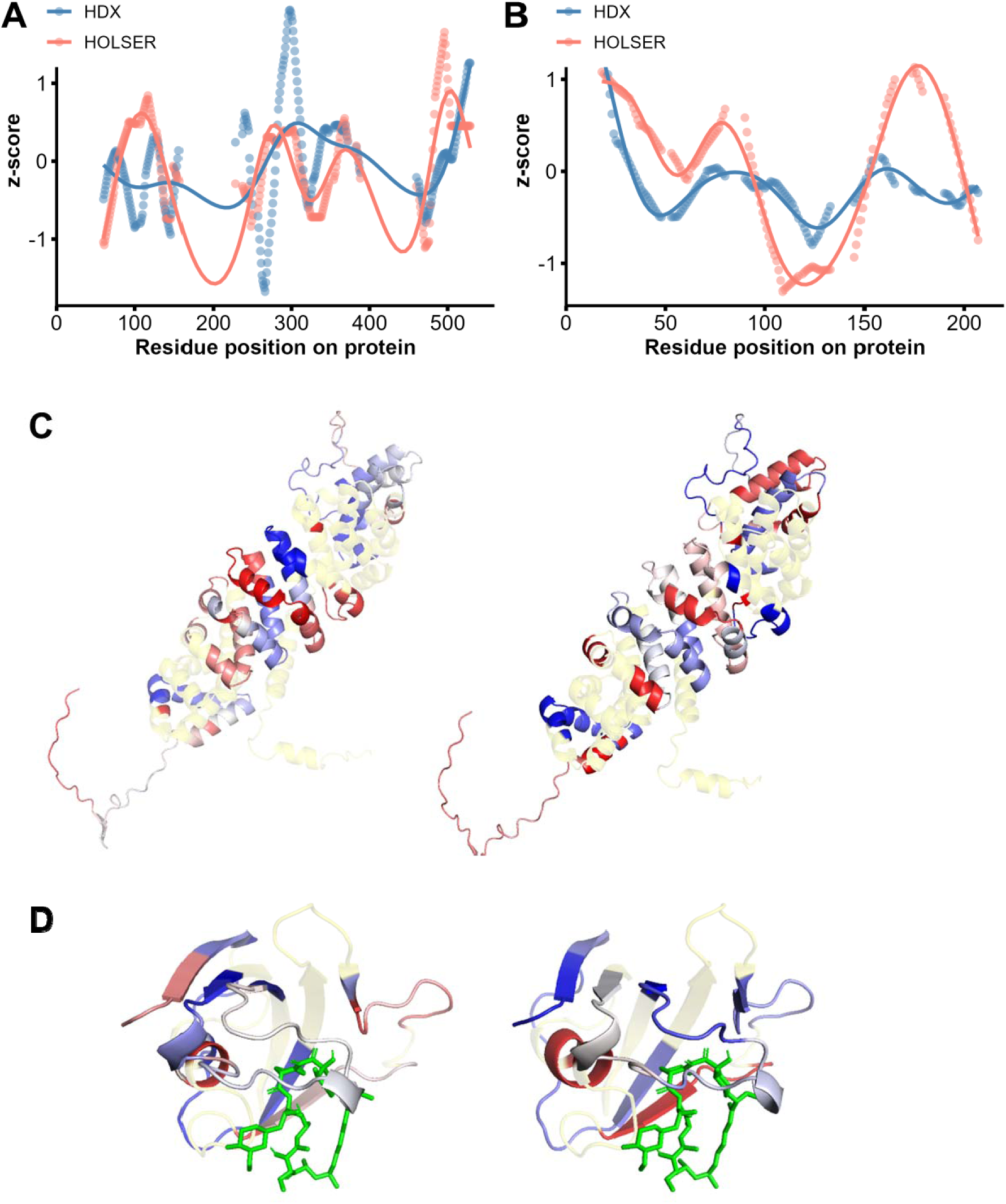
Analysis of accessibility. Residue-level accessibility comparison between HDX and HOLSER conditions for KPNA2 (**A**) and FKBP3 (**B**). Per-residue values were smoothed using a sliding window of 20 residues; each dot represents the average within that window. Smooth curves were fitted using a generalized additive model (GAM) with thin plate regression splines to highlight overall trends across the protein sequence. Protein structure (AF-P52292-F1-model_v4) for KPNA2 (**C**) and 1PBK for FKBP3 (**D**) colored by HDX MS (left) and HOLSER (right) z-scores in PyMOL. Blue means low accessibility, red means high accessibility, and light-yellow marks residues without data.

## CONCLUSIONS

We developed HOLSER, a partial proteolysis-based proteomics workflow that enables enhanced depth of proteome analysis, sequence coverage, improved quantification in detecting ligand-induced protein accessibility changes and provides global profiling of protein quaternary structures. The technique combines short digestion under native conditions with a high enzyme-to-substrate ratio, TMT-based multiplexing, and the inclusion of full-digest carrier channels. Compared to other existing approaches, HOLSER detects a greater number of peptides and proteins, yields significantly higher sequence coverage, and offers higher certainty in determining the binding site. Thus, HOLSER provides a robust and sensitive platform for profiling protein–ligand interactions under native conditions, with broad potential applications in target engagement studies, drug mechanism-of-action analysis, and proteome-wide conformational profiling.

## METHODS

### Cell culture

HeLa cells were purchased from ATCC^®^. Cells were grown at 37 LJ in Dulbecco’s Modified Eagle’s Medium (DEME, Thermo Fisher Scientific, Cat. 11594446) supplemented with 10% heat-inactivated fetal bovine serum (FBS, Thermo Fisher Scientific, Cat. 11560636) and 1% penicillin/streptomycin (Gibco, Cat. 15140-122) in a humidified atmosphere containing 5% CO_2_.

### HOLSER assay

Hela cells were collected and washed with PBS twice. The cells were resuspended in 20 mM EPPS (Merck, E9502) buffer at pH 8.2 containing protease inhibitors (Thermo Fisher Scientific, Cat. 78439) and were subjected to three freeze/thaw cycles to lyse the cells. The protein concentration was measured using a rapid protein quantification assay kit (Thermo Fisher Scientific, Pierce 660 nm) according to the manufacturer’s protocol. Then the lysate was diluted to 1 mg/mL with the lysis buffer (EPPS buffer at pH 8.2). 50 μL of cell lysates were incubated with 5 μL of the ligand solution in DMSO or DMSO only for 30 min at 25 °C in three replicates. The final concentration of the ligand was 10 μM for MTX, 10 μM for staurosporine and 2 μM for rapamycin. Limited trypsinolysis was carried out at an enzyme-to-substrate ratio of 1:1 (wt/wt) for a total duration of 4 min. Specifically, 10 µL of trypsin (Sigma-Aldrich, T1426) 5 mg/mL stock solution was added to 50 µg of protein sample. The mixture was gently and thoroughly mixed for 30 s using a multichannel pipette to ensure even distribution. The samples were then incubated at 25 °C for 3.5 min in ThermoMixer F1.5 (Eppendorf), being shaken at 1,000 rpm. For carrier proteome samples, trypsin digestion was carried out for 8 h. Proteolysis was stopped by adding 165 µL of 8 M guanidine hydrochloride (GdmCl) in 20 mM EPPS buffer (pHLJ8.2), resulting in a final concentration of 6 M GdmCl. To alkylate free cysteine residues via carbamidomethylation, tris(2-carboxyethyl)phosphine (TCEP) was added to a final concentration of 10 mM, followed by incubation at 95 °C for 5 min. Subsequently, iodoacetamide (IAA) was added to a final concentration of 40 mM. The samples were then diluted to a final concentration of 1 M guanidine hydrochloride (GdmCl) using 20 mM EPPS buffer (pH 8.2). The peptide samples were labelled using 16 TMTpro reagents (Thermo Fisher Scientific, Cat. A44520) according to the manufacturer’s instructions. The labelling reaction was quenched by 0.5% hydroxylamine (Thermo Fisher Scientific, Cat. 15225753). After pooling together the individually labeled samples, the peptides were acidified by trifluoroacetic acid (TFA) to a final concentration of 1% and desalted using C18 Sep-Pak cartridge. The cartridge was first conditioned by washing twice with 1 mL of 100% acetonitrile (ACN), followed by two washes with 1 mL of 0.1% TFA to equilibrate the resin. Acidified peptide samples were then loaded onto the cartridge and washed twice using 5% ACN with 0.1% formic acid (FA). Peptides were eluted from the cartridge using 50% ACN with 0.1% FA in two steps of 300 µL each. The eluates were collected and subsequently dried. Prior to LC-MS/MS analysis, dried peptides were reconstituted and fractionated by high-performance liquid chromatography with a Dionex Ultimate 3000 UPLC system (Thermo Fisher Scientific) as previously described^30, 31^. The generated 96 fractions were concatenated into 48 by pooling together every fourth fraction. For LC-MS/MS analysis, 1 µg of each pooled fraction was injected and analyzed.

### NanoLC-MS/MS analysis

NanoLC-MS/MS analyses were performed on an Orbitrap Exploris 480 mass spectrometer equipped with an EASY electrospray ionization source and connected online with an Ultimate 3000 nanoflow UPLC system (all - Thermo Fisher Scientific). The samples were pre-concentrated and desalted online using a PepMap C18 nano-trap column (length - 2 cm; inner diameter - 75 µm; particle size - 3 µm; pore size - 100 Å; Thermo Fisher Scientific) with a flow rate of 3 µL/min for 5 min. Peptide separation was performed on an EASY-Spray C18 reversed-phase nano-LC column (Acclaim PepMap RSLC; length - 50 cm; inner diameter - 2 µm; particle size - 2 µm; pore size – 100 Å; Thermo Fisher Scientific) at 55 °C and a flow rate of 300 nL/min. Peptides were separated using a binary solvent system consisting of 0.1% (v/v) FA and 2% (v/v) ACN (solvent A) as well as 98% ACN (v/v) and 0.1% (v/v) FA (solvent B). The peptides were eluted with a gradient of 3–26% B in 97 min, and 26–95% B in 9 min. Subsequently, the analytical column was washed with 95% B for 5 min before re-equilibration with 3% B. The mass spectrometer was operated in a data-dependent acquisition (DDA) mode. A survey mass spectrum (from m/z 375 to 1500) was acquired in the Orbitrap analyzer at a nominal resolution of 120,000. The automatic gain control (AGC) target was set as 100% standard, with the maximum injection time of 50 ms. In a 3 s DDA cycle, the most abundant ions with charge states 2^+^ to 7^+^ were isolated, fragmented using HCD MS/MS with 33% normalized collision energy, and detected in the Orbitrap analyzer at a nominal mass resolution of 50,000. The AGC target for MS/MS was set as 250% standard with a maximum injection time of 100 ms, whereas dynamic exclusion was set to 45 s with a 10-ppm mass window.

### HDX MS assay

Recombinant KPNA2 was kindly provided by Saei’s lab at Karolinska Institutet. Recombinant FKBP3 protein was obtained from Elabscience (PKSH032876). For HDX labeling, 4 μL of KPNA2 (5 μM) or FKBP3 (10 μM) was mixed with 36 μL of stock buffer prepared in DLJO (Sigma, 151882) and incubated at 10 °C for 30 s. The buffer composition for KPNA2 was 20 mM Tris-HCl, 50 mM NaCl, 2.5 mM DTT, pH 8.0, whereas for FKBP3 it was 20 mM Tris-HCl, 1 mM DTT, pH 8.0. After adding 1 volume of ice-cold 1.8M guanidine hydrochloride in 0.8% formic acid, the samples were analyzed in an automated HDX-MS system (TRAJAN) in which injected samples were automatically digested, cleaned and separated at 4 °C. Samples were digested using enzymeate BEH pepsin column (WATERS) followed by a 3-min desalting step using 0.1% FA at 100 µL/min. Peptic peptides were then separated by a 21 mm I.D × 50-mm length C18/3 µm column using a 21 min/10%–90% acetonitrile gradient in 0.1% formic acid at 40 µL/min. An Orbitrap Fusion Lumos mass spectrometer (Thermo Fisher Scientific) operated at 60,000 resolution at m/z 200 was used for analysis. Raw files were searched against the protein databases containing KPNA2 or FKBP3 aminoacidic sequence using MASCOT search engine. The HDExaminer software (Sierra Analytics) was used to process all HDX-MS data.

### Proteomics data analysis

LC-MS/MS data were analyzed using Proteome Discoverer version 3.1 (Thermo Scientific). Database searches were performed against the UniProt human reference proteome (Proteome ID: UP000005640_9606; downloaded in May 2025), containing 20,663 reviewed protein entries, for peptide and protein identification and quantification. TMT reporter ion abundances were normalized on the total abundance in a given TMT channel.

### Statistical analysis

Statistical significance of peptide abundance changes between treated and untreated samples was assessed using two-sided, unpaired Welch’s t-test, with p-values (p_p_) calculated across biological triplicates.

The peptide data were translated to the protein level by selecting top three peptides based on a combined score calculated as the absolute value of the log2 fold change in treated sample versus DMSO control multiplied by -log10(p_p_). The fold change for the protein was calculated as the average of those for the selected three peptides, and the protein p-value was obtained from the p_p_ values by combining them using Fisher’s formula. All statistical analyses were conducted using R (version 4.5.0). A p-value threshold of < 0.05 was considered statistically significant.

### Ligand–protein interaction mapping

The crystal structures of the studied protein–ligand complexes were retrieved and visualized in PyMOL (Schrödinger, LLC). The ligand and interacting residues were both shown in stick representation, with the ligand colored brown and the residues colored cyan. An embedded Python script was used to identify residues within the interaction interface. The script generated a tab-delimited file listing the interacting residues together with their chain ID, residue number, interaction type, and distance to the ligand.

### Accessibility analysis

Peptides with a coefficient of variation (CV) below 30% were selected for accessibility analysis. For HDX-MS, per-residue deuterium uptake was calculated by averaging the uptake values of all peptides covering each residue after 30 seconds of exchange. For HOLSER, the log2 fold change in peptide abundance between 4-minute and 8-hour trypsin digestions was assigned to each residue covered by the corresponding peptide. Both deuterium uptake and log2 fold change values were standardized across the protein using z-score transformation to enable direct comparison of relative accessibility. Z-score transformation was applied by subtracting the mean and dividing by the standard deviation of each dataset, centering the data and expressing each value in terms of standard deviations from the mean. To reduce local variability and capture broader structural trends, residues were grouped into sequential bins of 8 residues. Within each bin, the average HDX MS and HOLSER z-scores were computed along with the corresponding residue range. These binned values were then used to perform a Pearson correlation analysis, quantifying the relationship between HDX-MS and HOLSER-derived accessibility along the protein sequence. The data were first smoothed using a 20-point moving window to reduce noise and smoothing line is generated using a GAM with a spline on the x-axis, which produces a flexible, non-linear curve that follows the trend of the data without assuming a specific parametric form. For visualization, per-residue z-scores were mapped onto the three-dimensional structure of the protein using PyMOL. For KPNA2, the AlphaFold-predicted model (e.g., AF-P52292-F1-model_v4.pdb) was used. For FKBP3, the crystal structure (PDB ID: 1pbk) was employed for mapping. A custom blue–white–red color ramp was applied, representing low (blue), intermediate (white), and high (red) accessibility. Residues with no data in either dataset (i.e., not common to both HDX-MS and HOLSER) were shown in light yellow.

## Data availability

Excel files containing the analyzed data are provided in Supplementary Materials. The mass spectrometry proteomics data have been deposited to the ProteomeXchange Consortium (http://proteomecentral.proteomexchange.org) via the PRIDE partner repository with the dataset identifier PXD069119.

## Acknowledgements

R.A.Z. acknowledges support from EU (grant 956314 to ALLODD consortium), the Swedish Research Council (2021-05223) and Cancerfonden (22 1967 Pj). A.A.S. was supported by a grant from the Swedish Research Council (2023-02692) and the Novo Nordisk Foundation (NNF25OC0100921). We thank Dr. Majed Modaresi for purification of KPNA2.

---

**Supplementary Figure 1.**
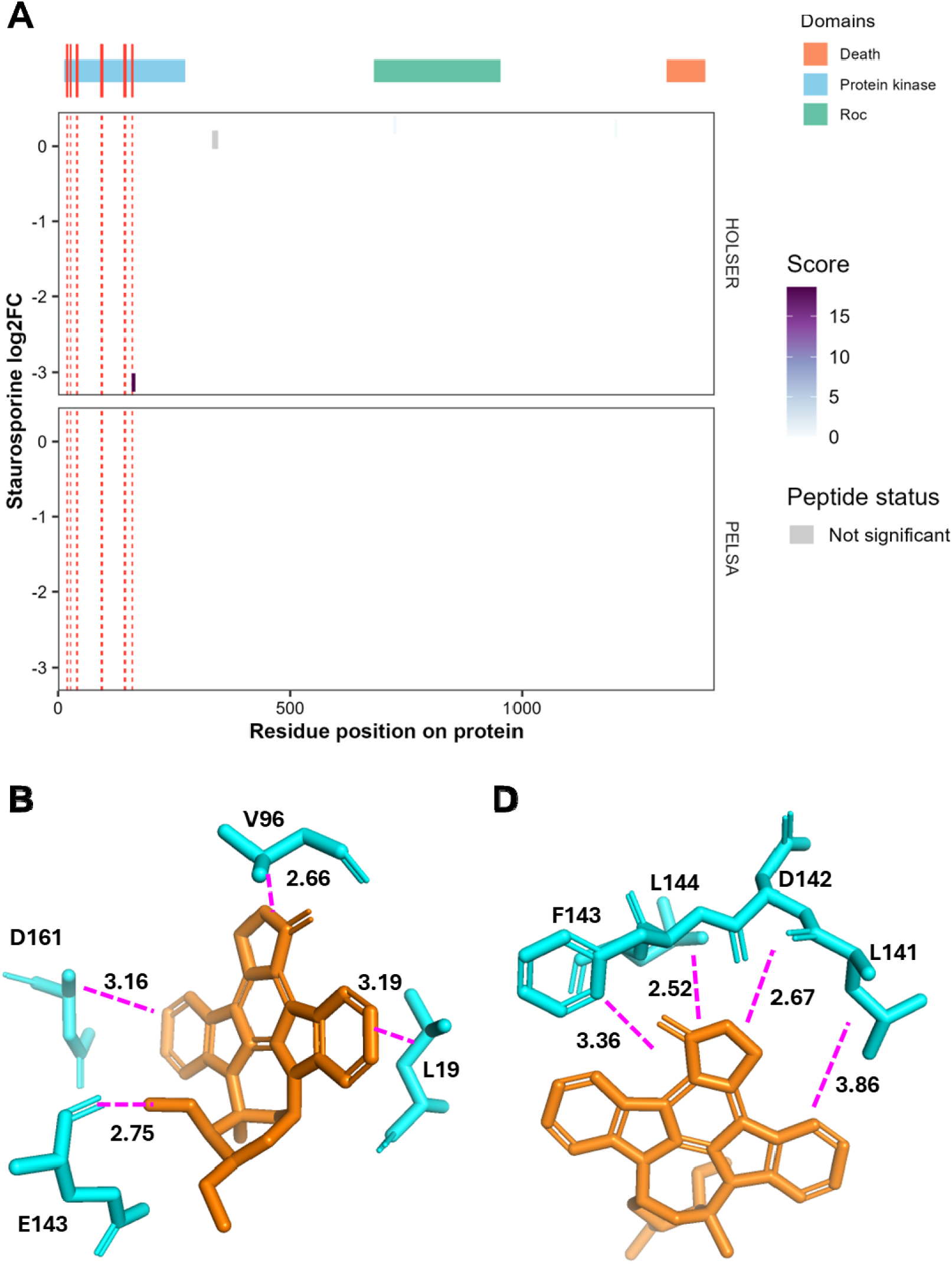
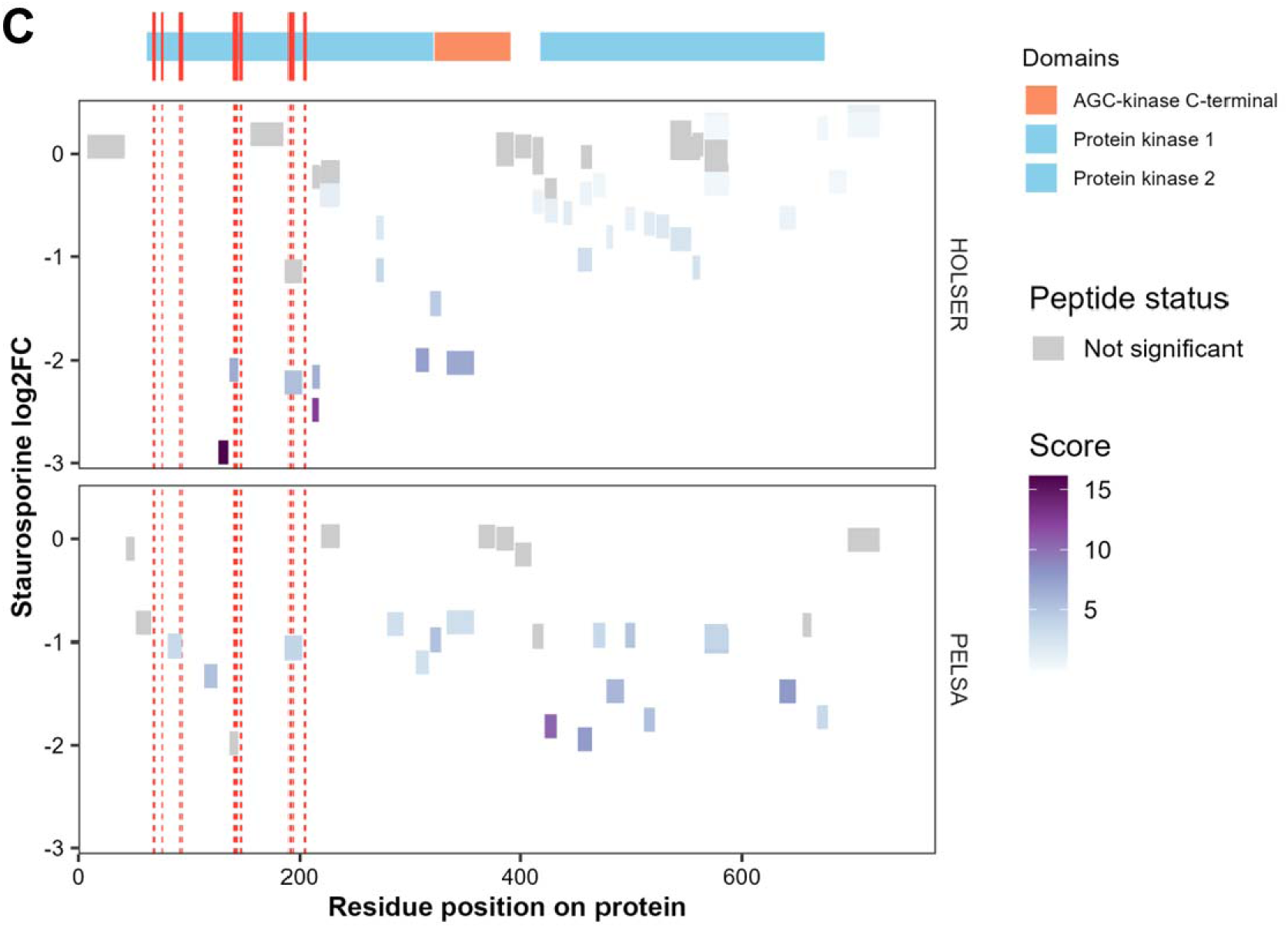
**A,** Mapping of DAKP1-derived peptides identified in HOLSER and PELSA onto the protein sequence. **B,** Structural visualization of the DAKP1–staurosporine complex based on the crystal structure (PDB ID: 1WVY), rendered in PyMOL. **C**, Mapping of RPS6KA1. **D**, Structural visualization of the RPS6KA1– staurosporine complex based on the crystal structure (PDB ID: 2Z7R), rendered in PyMOL.

